# Yersiniabactin producing AIEC promote inflammation-associated fibrosis in gnotobiotic *Il10^−/−^* mice

**DOI:** 10.1101/723148

**Authors:** Melissa Ellermann, Raad Z Gharaibeh, Laura Fulbright, Belgin Dogan, Lyndsey N Moore, Christopher A. Broberg, Lacey R. Lopez, Aaron M. Rothemich, Jeremy W Herzog, Allison Rogala, Ilyssa O. Gordon, Florian Rieder, Cory R. Brouwer, Kenneth W. Simpson, Christian Jobin, R Balfour Sartor, Janelle C Arthur

**Affiliations:** Department of Microbiology and Immunology, University of North Carolina at Chapel Hill, NC, USA; Center for Gastrointestinal Biology and Disease, University of North Carolina at Chapel Hill, NC, USA; Lineberger Comprehensive Cancer Center, University of North Carolina at Chapel Hill, NC, USA; Department of Medicine, University of Florida, Gainesville, FL, USA; Department of Infectious Diseases and Pathology, University of Florida, Gainesville, FL, USA; Department of Clinical Sciences, College of Veterinary Medicine, Cornell University, Ithaca, NY, USA; Department of Pathology, Robert J. Tomsich Pathology and Laboratory Medicine Institute, Cleveland Clinic Foundation, Cleveland, USA; Department of Gastroenterology, Hepatology and Nutrition, Digestive Diseases and Surgery Institute, Cleveland Clinic Foundation, Cleveland, USA; Department of Inflammation and Immunity, Lerner Research Institute, Cleveland Clinic Foundation, Cleveland, USA; Department of Bioinformatics and Genomics, University of North Carolina at Charlotte, Charlotte, NC, USA

**Keywords:** Fibrosis, AIEC, Crohn’s disease, colitis, microbiome, yersiniabactin

## Abstract

Fibrosis is a significant complication of intestinal disorders associated with microbial dysbiosis and pathobiont expansion, notably Crohn’s disease (CD). Mechanisms that favor fibrosis are not well understood and therapeutic strategies are limited. Here we demonstrate that colitis susceptible *Il10*-deficient mice develop inflammation-associated fibrosis when mono-associated with adherent/invasive *Escherichia coli* (AIEC) that harbor the yersiniabactin (Ybt) pathogenicity island. Inactivation of Ybt siderophore production in AIEC nearly abrogated fibrosis development in inflamed mice. In contrast, inactivation of Ybt import through its cognate receptor FyuA enhanced fibrosis severity. This corresponded with increased colonic expression of profibrogenic genes prior to the development of histological disease, therefore suggesting causality. *FyuA*-deficient AIEC also exhibited greater localization within sub-epithelial tissues and fibrotic lesions that was dependent on Ybt biosynthesis and corresponded with increased fibroblast activation *in vitro*. Together, these findings suggest that Ybt establishes a pro-fibrotic environment in the host in the absence of binding to its cognate receptor and indicates a direct link between intestinal AIEC and the induction of inflammation-associated fibrosis.

## Introduction

Inflammatory bowel diseases (IBD), including Crohn’s disease (CD), are characterized by chronic intestinal inflammation that develops as a result of prolonged and inappropriate mucosal immune responses to luminal antigens in genetically susceptible individuals (1). The chronic and relapsing nature of IBD, in conjunction with the lack of curative therapies for many patients, enhances risk for inflammation-associated comorbidities including intestinal fibrosis (2). Approximately 30% of CD patients develop fibrotic disease that can result in intestinal strictures and bowel obstructions (2) (3) (4). Current treatments for intestinal fibrosis are inadequate and rely on anti-inflammatory therapies (which are often ineffective) and surgical interventions (3). Fibrosis is recurrent in large proportions of the CD population (4), thus necessitating the development of specific anti-fibrotic therapeutics.

Fibrosis is characterized by excess accumulation of extracellular matrix (ECM) components that results in the pathological remodeling of tissues and consequent organ dysfunction. Mesenchymal cells such as fibroblasts, myofibroblasts and smooth muscle cells become highly activated in response to transmural injury or inflammation and hypersecrete ECM components and profibrogenic factors that further propagate fibrotic processes. The tissue microenvironment also plays an important role in modulating the activity of mesenchymal cells, where host-derived signals such as cytokines and growth factors serve as additional fibrogenic or antifibrotic mediators (4) (3). Activation of mesenchymal cells is also subject to regulation by microbial factors (5) (6). Fibrosis can occur in bacterial-induced models of acute colitis including mice chronically colonized with the enteric pathogen *Salmonella enterica* or with a CD-associated *Escherichia coli* pathobiont (7) (8). Importantly, progression from intestinal inflammation to inflammation-associated fibrosis is incompletely penetrant in bacterial-induced colitis models and in clinical populations with microbial-driven diseases like IBD. It remains unclear which microbiota-derived signals favor the establishment of a profibrogenic microenvironment.

The intestinal microbiota are key modulators of mucosal immunity under homeostatic conditions and in numerous inflammatory pathologies including IBD (1). A subset of resident intestinal *E. coli* known as adherent and invasive *E. coli* (AIEC) are enriched in CD patients (9) (10) (11). AIEC breach the intestinal epithelium and induce inflammation in various rodent models of experimental colitis (12) (13) (14) (15). Colonization of germ free, inflammation-prone *Il10^−/−^* mice with AIEC induces aggressive, transmural intestinal inflammation driven by bacterial antigen-specific T-helper-(Th)1 and Th17 immune responses (13) (16). Studies in germ free *Il10^−/−^* mice individually colonized with AIEC have led to the identification of several bacterial factors that augment or diminish the colitis-inducing and pro-carcinogenic capabilities of AIEC (17) (18) (19) (20).

Comparative phylogenetic studies have demonstrated that the yersiniabactin (Ybt) high pathogenicity island (HPI) is overrepresented in human, canine and murine AIEC strains (21). The Ybt HPI encodes enzymatic machinery required for the biosynthesis of the siderophore Ybt (22). Once Ybt is released from bacterial cells, it sequesters extracellular metals including iron, zinc and copper. The Ybt-metal chelate is subsequently imported through its cognate outer membrane receptor FyuA for bacterial use (22) (23) (24). The Ybt HPI is harbored by numerous Enterobacteriaceae pathogens and contributes to *in vivo* fitness, niche formation, and virulence (25) (26) (27). However, the contribution of the Ybt HPI to the proinflammatory potential of resident intestinal *E. coli* such as AIEC has not been explored, despite its prevalence in this population. We therefore utilized our gnotobiotic *Il10^−/−^* mouse model to investigate whether inactivation of the Ybt system in AIEC modulates immune-mediated colitis. While abrogation of Ybt biosynthesis in AIEC delayed colitis onset, colonization of mice with Ybt-positive AIEC was associated with the development of inflammation-associated fibrosis. Severity of fibrosis was enhanced in mice colonized with the Ybt-positive transport mutant (Δ*fyuA*), which corresponded with increased profibrogenic gene signatures in the colon and in cultured fibroblasts and enhanced AIEC subepithelial localization within fibrotic lesions. Abrogation of Ybt biosynthesis in Δ*fyuA* attenuated fibrosis in inflamed mice, restored AIEC localization to the epithelium and reduced fibroblast activation. Collectively, our findings introduce a non-canonical role for Ybt in mediating fibrosis development independent of its established function in delivering iron to bacteria through FyuA. More broadly, we introduce a novel microbial-driven, immune-mediated model of inflammation-associated fibrosis that recapitulates key histopathological features of fibrotic disease in human CD.

## Results

### Inactivation of Ybt biosynthesis, but not Ybt transport, in AIEC delays progression of colitis

The siderophore Ybt and its cognate receptor FyuA mediate bacterial metal acquisition in pathogenic Enterobacteriaceae. Because the Ybt HPI is also harbored by many IBD-associated AIEC strains, we hypothesized that like its pathogenic counterparts, the Ybt HPI enhances the proinflammatory potential of AIEC. To determine whether an intact Ybt siderophore system in AIEC contributes to colitis development, we inactivated Ybt biosynthesis or import by creating isogenic mutants unable to import Ybt-metal chelates (Δ*fyuA)* or unable to synthesize Ybt *(Δirp1)* in the AIEC strain NC101 (which also harbors the enterobactin and salmochelin siderophore systems). We colonized germ-free, inflammation-susceptible *Il10^−/−^* mice with NC101, Δ*fyuA* or Δ*irp1* and compared the severity of colitis induction. At 5 weeks, colitis histopathology was significantly attenuated in mice colonized with Δ*irp1* compared with Ybt+ NC101 and Δ*fyuA* (Fig. 1A-E) an attenuation that was no longer apparent by 10 weeks (Fig. S1). In contrast, colitis development did not differ in mice colonized with NC101 versus Δ*fyuA*. Colitis scores differences did not correlate with altered expression of proinflammatory cytokines known to correlate with disease in this model (Fig. S1) (13) (16). The reduced colitis potential of Δ*irp1* did not correspond with diminished luminal growth in the gut (Fig. 1G-I) or *in vitro* growth defects under iron replete or limiting conditions (Fig. S2). While the Δ*fyuA* mutant exhibited a growth defect at 5 weeks, its attenuated growth was not sustained throughout colitis development and did not correlate with colitis severity (Fig. 1G-I). Together, these findings demonstrate that Ybt enhances the proinflammatory potential of AIEC in gnotobiotc, inflammation-susceptible hosts.

**Figure 1.**
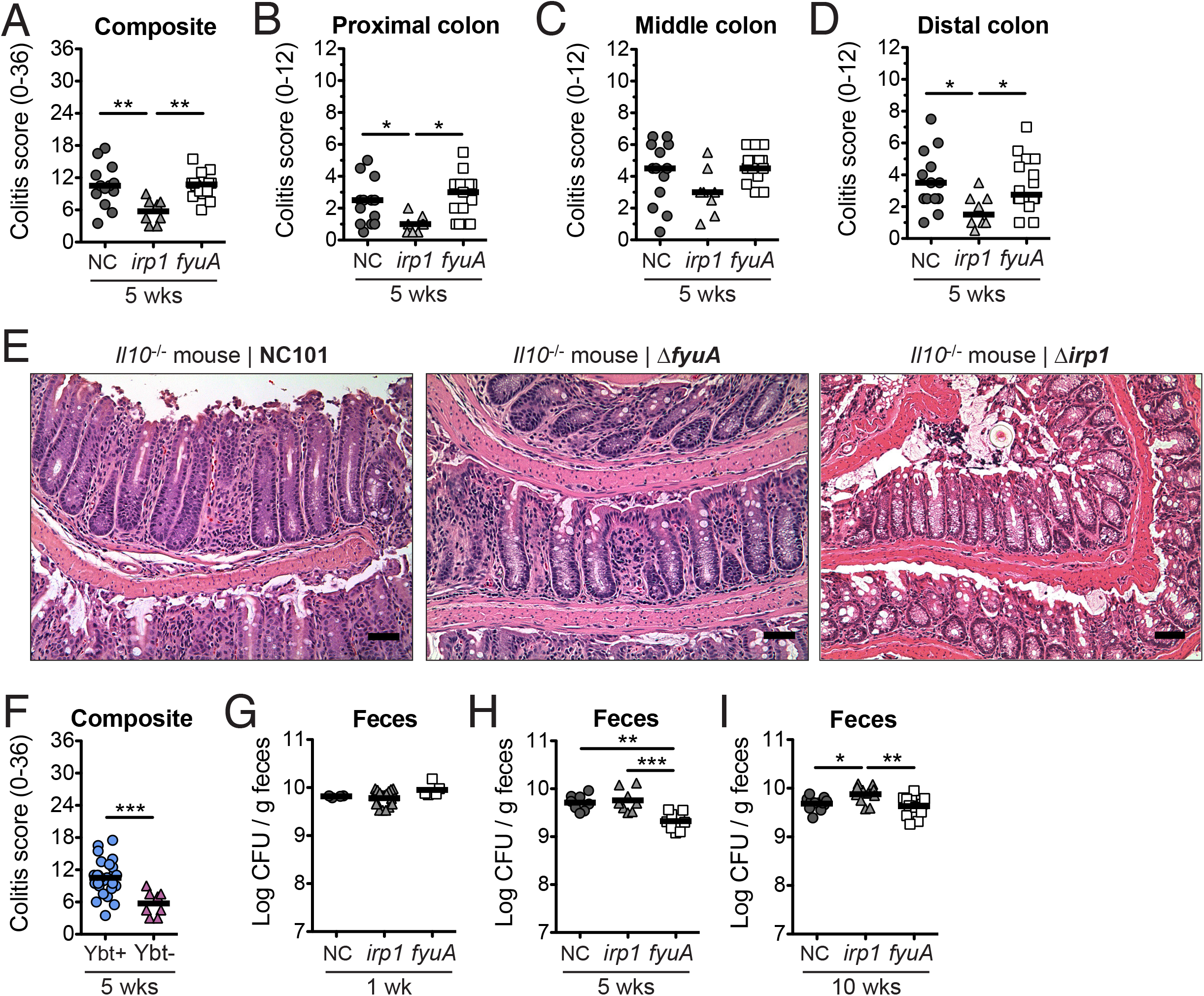
Yersiniabactin enhances the proinflammatory potential of AIEC in gnotobiotic *Il10^−/−^* mice. Germ free *Il10*^−/−^ mice were mono-associated with the AIEC strain *E. coli* NC101 (NC), Δ*fyuA* or Δ*irp1* for 5 weeks. A) Composite and B-D) regional histopathology colitis scores. E) Representative H&E histology of the colon. Scale bar, 50 μm. F) Composite histopathology colitis scores of *Il10^−/−^* mice colonized with yersiniabactin-(Ybt)-positive or Ybt-deficient NC101. Lines are at the median. *P*-values were determined by Kruskal-Wallis or Mann-Whitney. G-I) Quantitative bacteria culture from feces at G) 1 week, H) 5 weeks or I) 10 weeks post-colonization. Lines are at the mean. *P*-values were determined by one-way ANOVA. Each symbol represents an individual mouse (n = 8-14). * *p* < 0.05, ** *p* < 0.01, *** *p* < 0.001.

### Ybt-positive AIEC promote fibrosis development in inflamed *Il10^−/−^* mice

In a subset of NC101- and Δ*fyuA*-colonized inflamed *Il10^−/−^* mice, but rarely in Δ*irp1*-colonized *Il10^−/−^* mice, pathological remodeling of the colonic submucosa was observed in hematoxylin and eosin (H&E) stained colon sections (Fig. 2, S3). Histological features consistent with fibrosis, including marked expansion of the submucosa with excessive deposition of lightly eosinophilic, fibrillar substances, characterized the pathology. Positive staining with Masson’s trichrome and Sirius red confirmed the presence of collagen fibers as part of the expanded ECM in fibrotic mice (Fig. 2B). Lamina propria collagen localization was also altered in fibrotic mice, exhibiting a basal predilection. In contrast, in non-fibrotic AIEC-colonized *Il10^−/−^* mice, the submucosal ECM was structured and organized and stained collagen fibrils in the lamina propria exhibited an apical propensity (Fig. 2A). Taken together, a subset of AIEC-colonized *Il10^−/−^* mice develop histopathological lesions that are consistent with fibrosis.

**Figure 2.**
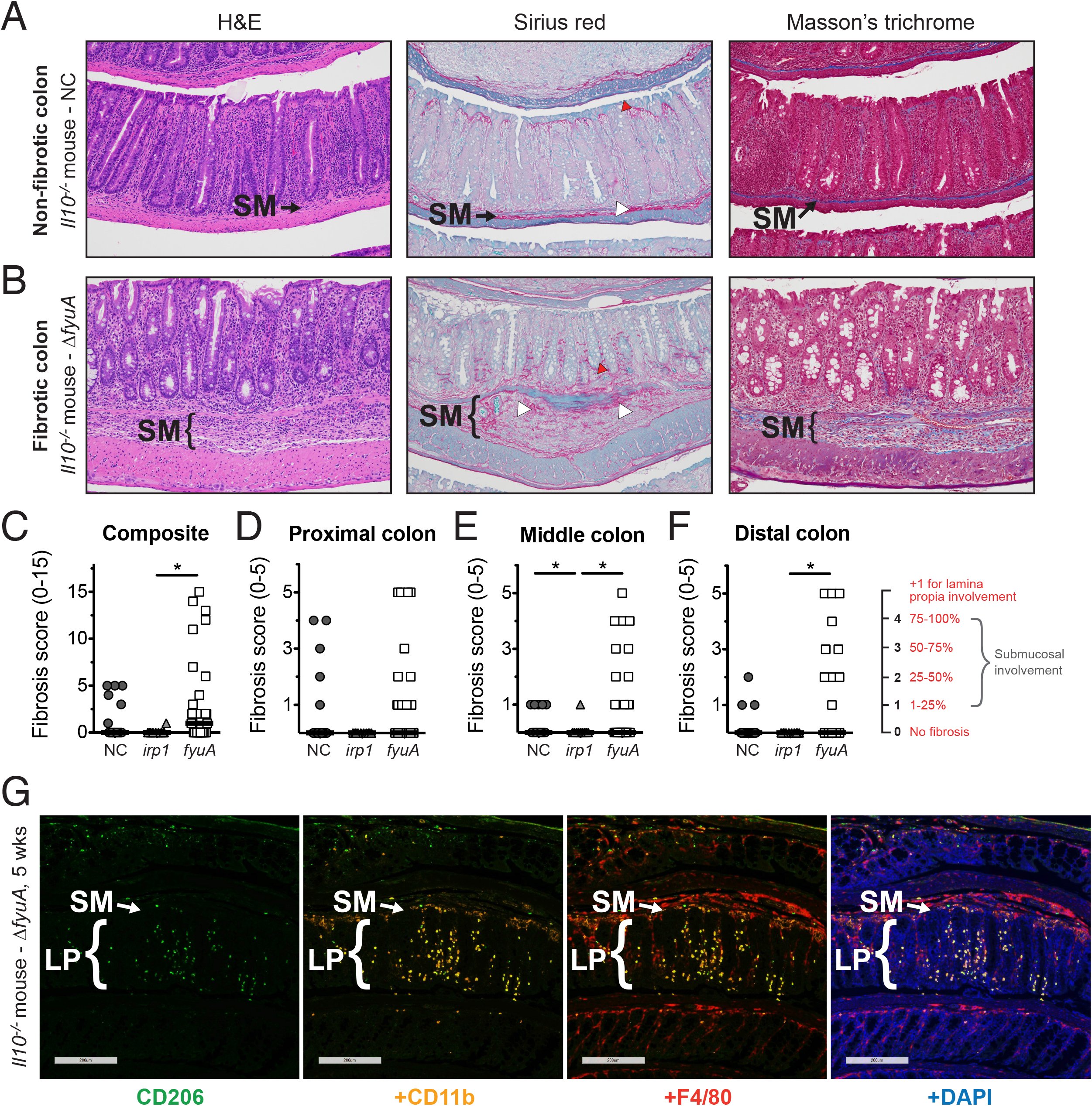
Ybt+ AIEC promotes fibrosis development in colitic *Il10^−/−^* mice. Germ free *Il10^−/−^* mice were mono-associated with the Ybt+ AIEC strains NC or Δ*fyuA* or the Ybt-strain Δ*irp1* for 10 weeks. A-B) Representative colonic histology of gnotobiotic *Il10^−/−^* mice colonized with A) NC or B) Δ*fyuA*. Colon sections were stained with H&E, Sirius red/fast green or Masson’s trichrome. Regions of Sirius red binding is indicated by white arrowheads in the submucosa and red arrowheads in the lamina propria. C) Composite and D-F) regional fibrosis histology scores. Each symbol represents an individual mouse (n = 11-29). Lines are at the median. *P*-values were determined by Kruskal-Wallis. * *p* < 0.05. G) Representative colonic histology from gnotobiotic *Il10^−/−^* mice colonized with Δ*fyuA* for 5 weeks. Colonic sections were stained with antibodies against the established macrophage cell surface markers CD206, CD11b, and F4/80 and were counterstained with the DNA stain DAPI. Scale bar, 200 μm. SM, submucosa. LP, lamina propria. L, lumen.

Because fibrosis incidence seemed to differ between NC101-, Δ*fyuA-, and Δirp1-* colonized *Il10^−/−^* mice, we next utilized a fibrosis pathology scoring system to determine whether the Ybt system in AIEC impacts inflammation-associated fibrosis (28) (29) (see Materials and Methods). The most severe fibrosis pathology in all regions of the colon were observed in Δ*fyuA*-colonized *Il10^−/−^* mice, which corresponded with higher incidence of severe disease. In contrast, moderate-severe fibrosis in NC101-colonized mice was mostly restricted to proximal colon (Fig. 2C-F). These differences in fibrosis severity and incidence were associated with altered cellular populations infiltrating the submucosa, with immunologically-defined macrophages (CD206+, CD11b+ and/or F4/80+ cells) observed in Δ*fyuA*-colonized fibrotic mice (Fig. 2G) versus the inflammatory lymphocytes consistently observed in NC101-colonized, non-fibrotic mice (20). Inflamed *Il10^−/−^* mice colonized with the Ybt-deficient Δ*irp1* mutant did not develop moderate-severe fibrotic lesions and rarely exhibited mild disease (Fig. 2A-D), suggesting a role for Ybt in inducing and exacerbating this pathology. To validate that the histopathology in our mouse model is consistent with inflammation-associated fibrosis in human CD, we evaluated H&E and Sirius Red staining of full-thickness colon resection tissues from fibrotic CD, ulcerative colitis, diverticulitis, and healthy margins of colorectal cancer resections (Fig. 3, S5). Fibrotic CD tissues exhibited remarkable similarity to our mouse model, with transmural inflammation, expansion of the submucosa, thick collagen fibrils, and disruption of the muscularis by collagen and infiltrating cells (Fig. 3). Fibrosis was not evident by H&E, Sirius Red, or Masson’s Trichrome or at 5 weeks in *Il10^−/−^* mice (data not shown). Collectively, these observations demonstrate that Ybt+ AIEC promote the development of fibrotic disease in an experimental model of pathobiont-induced colitis.

**Figure 3.**
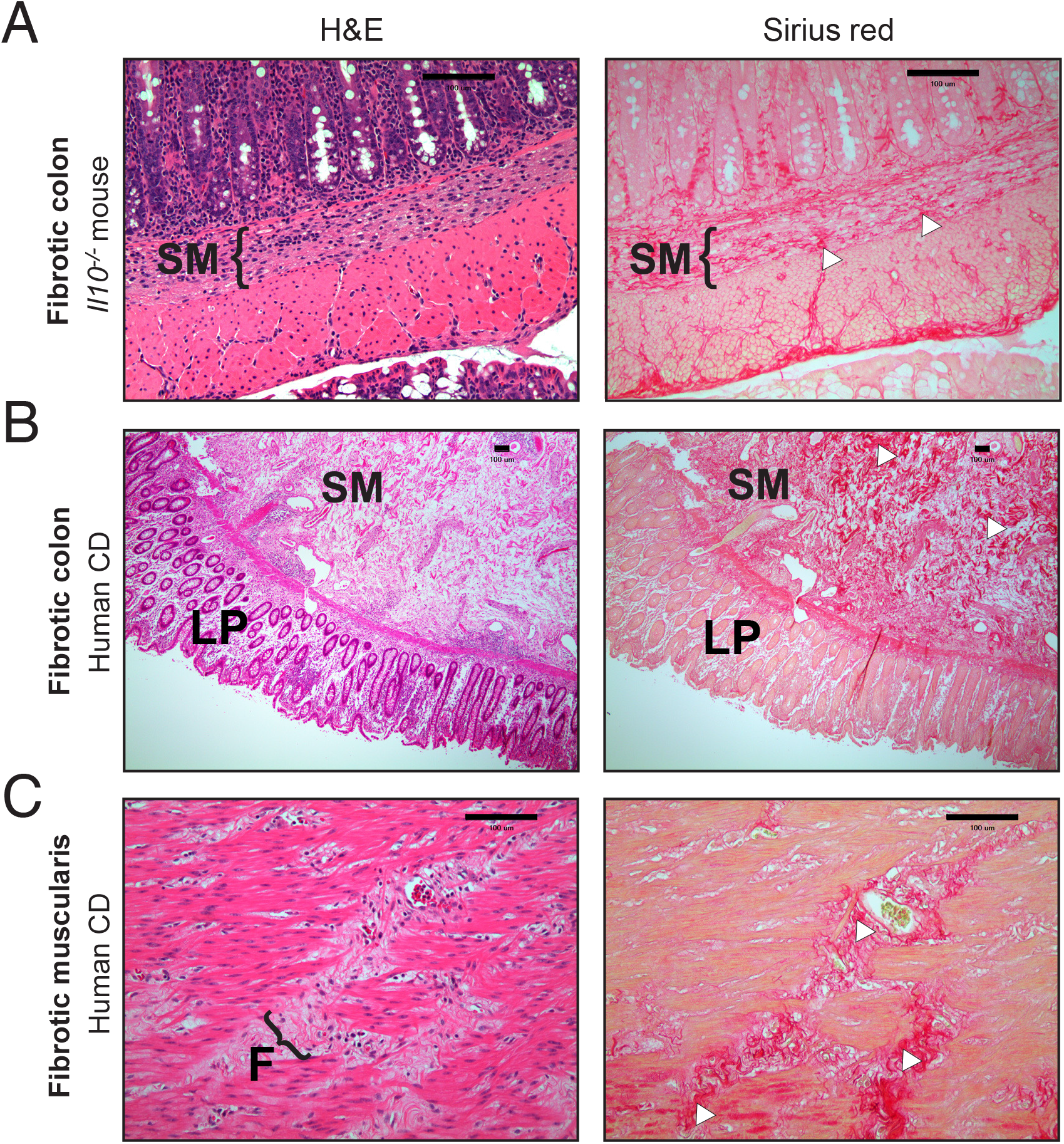
Fibrosis development in AIEC-colonized *Ill10^−/−^* mice recapitulates histopathological features of fibrosis in human Crohn’s disease. Representative colonic histology of A) Δ*fyuA*-colonized fibrotic *Il10^−/−^* mice. B-C) Representative histology of full thickness colon cross-sections from fibrotic Crohn’s disease patients, representative of *n* = 3 per group. C) Magnification of the muscularis serosa. Colon sections were stained with H&E or Sirius red. Regions of Sirius red binding are indicated with white arrowheads. LP, lamina propria. SM, submucosa. F, fibrotic lesion. Scale bar, 100 μm.

Because fibrosis occurs in response to tissue injury instigated by inflammation, we next determined whether fibrosis severity positively correlates with inflammation. Linear regression analysis revealed a significant negative correlation between fibrosis and colitis histopathology in the middle colon and no significant correlations in the proximal and distal colon (Fig. S4). Moreover, NC101- and Δ*fyuA*-colonized mice exhibited similar levels of colitis histopathology despite the exacerbated fibrosis observed in Δ*fyuA*-colonized mice (Fig. 1–2). Nonetheless, as previously reported (4), inflammation is required for the pro-fibrotic activities of NC101 and Δ*fyuA* given that fibrosis was not observed in uninflamed WT mice colonized with either strain (Fig. S3). These results demonstrate that while inflammation is required for fibrosis development, Ybt+ AIEC exacerbate inflammation-associated fibrosis independent of effects on the proinflammatory potential of AIEC.

### Fibrosis development corresponds with enhanced subepithelial invasion of *fyuA*-deficient AIEC

We next determined whether the pro-fibrogenic potential of Ybt+ AIEC corresponds with altered bacterial localization within the intestines. While colonic mucus colonization did not differ between the strains, colonic tissue loads of AIEC were significantly increased in Δ*fyuA*-colonized mice at 10 weeks (Fig. 4A-B). In contrast, colonic tissue colonization did not differ between NC101 and Δ*irp1*. Colonic mucus or tissue loads were also comparable at 5 weeks (Fig. S6A-B). Because AIEC are functionally characterized by epithelial invasiveness, intra-macrophagic survival and robust biofilm formation, we performed standard *in vitro* assays commonly utilized to distinguish AIEC strains (9). While iron availability altered AIEC epithelial invasion, no differences in epithelial adherence or invasion were observed between NC101, Δ*irp1* or Δ*fyuA* under iron replete or limiting conditions (Fig. S6C-D). Similarly, genetic ablation of Ybt transport did not alter macrophage phagocytosis or intracellular survival of AIEC (Fig. S6E-G) and had no effect on AIEC biofilm formation (Fig. S6H). Thus, while Ybt transport or biosynthesis did not impact defining *in vitro* characteristics of AIEC, deletion of *fyuA* enhanced AIEC colonic tissue colonization, suggesting that FyuA may be important in modulating bacterial localization within the intestines.

**Figure 4.**
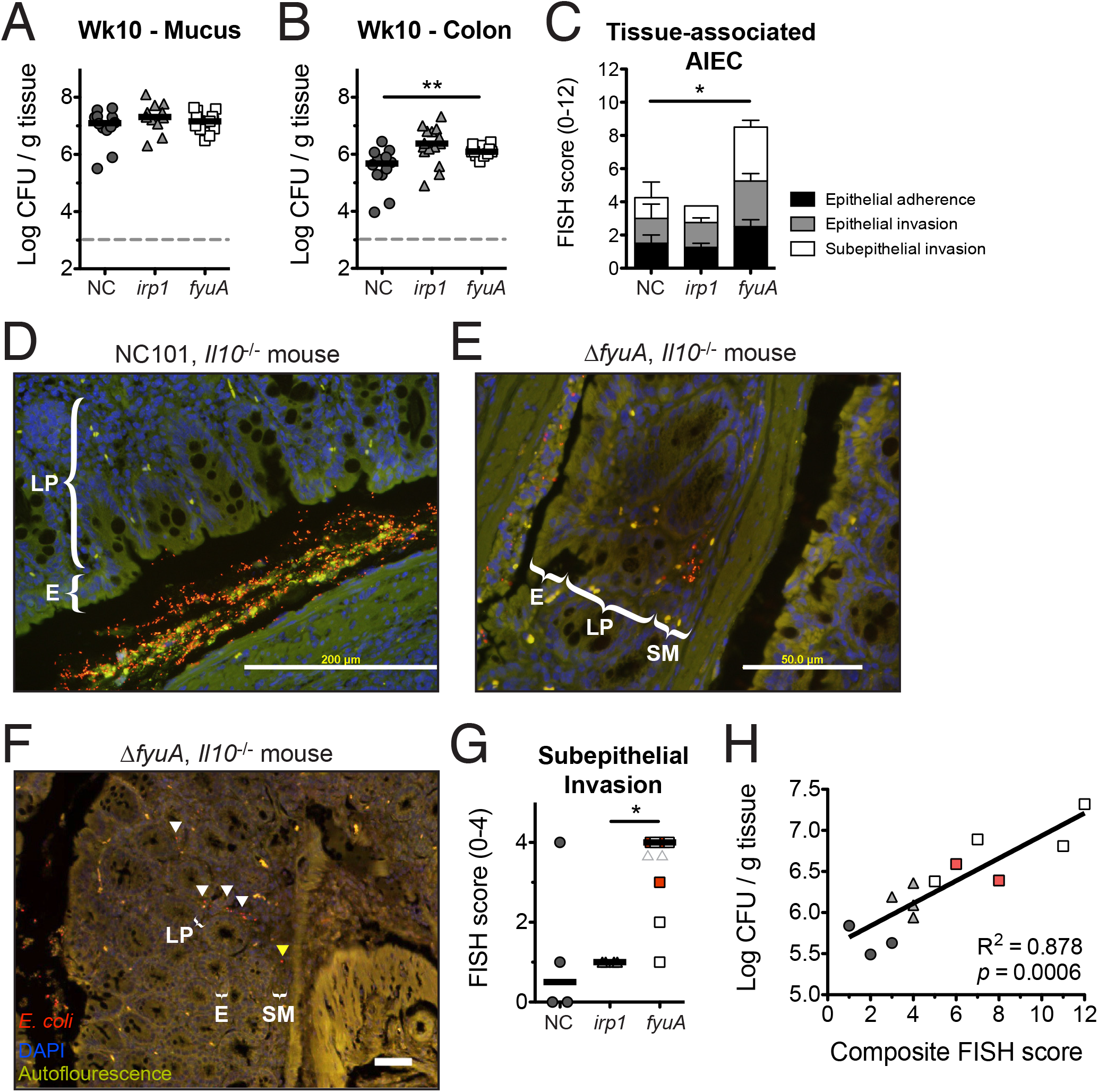
Inactivation of yersiniabactin transport enhances AIEC mucosal invasion. Germ free *Il10^−/−^* mice were mono-associated with NC, Δ*fyuA* or Δ*irp1* for 10 weeks. Quantitative bacterial culture of A) colonic mucus or B) colonic tissues. Each symbol represents an individual mouse (n = 11-15). Lines are at the median. *P*-values were determined by Kruskal-Wallis. C) FISH analysis of proximal colons (*n* = 4-8). *P*-values were determined by Kruskal-Wallis. D-F) Representative FISH images of the proximal colon. Red, *E. coli*. Blue, DAPI. Arrowheads in F) indicate *E. coli* localized within the lamina propria (white) and submucosa (yellow). E, epithelium. LP, lamina propria. SM, submucosa. Scale bar, 200 μm. G) Subepithelial AIEC invasion scores as assessed by FISH in C. Red squares (indicated by the grey triangles) represent fibrotic mice as assessed by histopathology. Lines are at the median. *P*-values were determined by Kruskal-Wallis. H) Linear regression analysis of quantitative bacterial culture versus FISH score from colonic tissues. Red squares represent fibrotic mice as assessed by histopathology. * *p* < 0.05, ** *p* < 0.01.

To further assess how FyuA impacts AIEC localization in the gut, we employed a more sensitive approach – *E. coli* 16S fluorescence in situ hybdrization (FISH) – to visualize tissue-associated AIEC. FISH analysis revealed an overall increase in tissue-associated Δ*fyuA* relative to NC101 and Δ*irp1* (Fig. 4C). This difference was primarly driven by enhanced subepithelial (lamina propria and submucosa) localization of Δ*fyuA* (Fig. 4D-G). Moreover, Δ*fyuA* was observed within submucosal fibrotic lesions, demonstrating its co-localization with diseased tissue (Fig. 4F, arrowheads). Importantly, tissue bacteria loads assessed by quantitative bacterial culture and FISH analysis were positively correlated (Fig. 4H). Together, these results suggest that inactivation of *fyuA* enhances the subepithelial localization of AIEC, which may contribute to its profibrogenic potential.

### Inactivation of Ybt-mediated metal acquisition does not alter AIEC iron sensing

The canonical function of Ybt is to scavenge extracellular metals for bacterial use (22) (31). Because the most severe fibrosis occurred in mice colonized with Δ*fyuA*, we first assessed whether Ybt functionality was altered in this mutant. The extent of Ybt secretion was comparable between NC101 and Δ*fyuA*, and as expected, Ybt secretion was not detected in Ybt biosynthesis mutant Δ*irp1* (Fig. S7a). We next confirmed the functionality of Ybt produced by NC101 and Δ*fyuA*. To accomplish this, we assessed whether Ybt produced by these strains can restore the growth of siderophore-deficient *Klebsiella pneumoniae* (Δ*entB irp1*) cultivated under iron-limiting conditions. In contrast to Δ*irp1*, both Ybt+ NC101 and Δ*fyuA* rescued *K. pneumoniae* Δ*entB irp1* growth (Fig. S7b). Taken together, these data suggest altered Ybt functionality does not correspond with the increased profibrogenic potential of Δ*fyuA*.

Mutants lacking FyuA are unable to import Ybt-iron chelates and may therefore be unable to satisfy their iron requirements. Thus, the enhanced profibrogenic potential of Δ*fyuA* may be the result of altered bacterial function mediated through disrupted bacterial iron homeostasis. To test this idea, we first compared *in vivo* expression of iron-responsive genes in NC101, Δ*fyuA* and Δ*irp1*. Transcript levels of several iron-responsive genes did not differ between strains (Fig. S8A-B), suggesting that NC101 iron homeostasis is not perturbed upon inactivation of Ybt transport or biosynthesis in the intestines. Similarly, *in vitro* iron depletion with the iron chelator 2’2,bipyridyl (BPD) did not alter transcription of iron responsive genes (Fig. S8C).

To corroborate these results, we performed transcriptional reporter assays utilizing vectors harboring *gfp* fused to the iron-responsive promoter P_*tonB*_. To first validate this approach, the NC101 reporter strain was cultivated under iron replete and limiting conditions, and as expected, iron depletion enhanced *gfp* activity driven by the *tonB* promoter (Fig. S9a). We next assessed whether NC101, Δ*fyuA* and Δ*irp1* iron-sensing reporters respond differently to iron depletion. In agreement with our transcriptional results, *gfp* expression was comparable between NC101 and Δ*fyuA* (Fig. S9a), suggesting that inactivation of FyuA does not impact AIEC iron sensing. Because Ybt can also bind other metals including zinc (our own observations) (22) (47) and copper (our own observations) (23) (24), we performed similar assays with the zinc responsive promoter P_*znuA*_ and the copper responsive promoter P_*cusC*_. As with the iron sensing reporters, altering zinc and copper availability did not alter sensing of the respective metals in Δ*fyuA* relative to NC101 (Fig. S9b-c). In contrast, the activities of iron- and zinc-responsive promoters were significantly increased in Δ*irp1* (Fig. S9a-b), suggesting that metal starvation is enhanced in this mutant under iron and zinc limiting conditions. Taken together, these data suggest that while metal sensing in AIEC is not altered with disruption of Ybt transport, metal homeostasis appears disrupted in Ybt-negative Δ*irp1*.

### Deletion of *fyuA* in AIEC promotes the establishment of a pro-fibrotic colonic environment that precedes fibrosis development

The increased incidence of fibrosis in Δ*fyuA*-colonized *Il10^−/−^* mice may in part be driven by differential host responses to fyuA-expressing versus fyuA-deficient AIEC. To test this idea, we utilized high-throughput RNA sequencing (RNAseq) to determine whether global differences are apparent in the colonic transcriptomes of inflamed *Il10^−/−^* mice and non-inflamed WT mice colonized with NC101 or Δ*fyuA*. Principal Coordinate Analysis (PCoA) revealed significant differences in the colonic transcriptomes of NC101-colonized non-fibrotic versus Δ*fyuA-* colonized fibrotic *Il10^−/−^* mice at 10 weeks when fibrosis is apparent (Fig. 5a). This corresponded with 2692 genes and 71 KEGG pathways that were differentially expressed between Δ*fyuA-* versus NC101-colonized *Il10^−/−^* mice (Table S3, S4). In contrast, the transcriptomes of NC101-versus Δ*fyuA-* colonized WT mice clustered together (Fig. 5a), suggesting that differences in the host transcriptional responses to either strain predominantly occur in *Il10^−/−^* mice. To determine whether the differing host responses precede histological evidence of fibrosis, we also compared the colonic transcriptomes of NC101-colonized versus Δ*fyuA-* colonized *Il10^−/−^* mice at 5 weeks. RNAseq analysis revealed that 169 genes and 116 KEGG pathways were differentially expressed in NC101-colonized versus Δ*fyuA-* colonized *Il10^−/−^* mice (Table S1, S2), many of which were differentially regulated at both 5 and 10 weeks. However, testing overall community composition did not reach statistical significance after FDR correction (p < 0.084) (Fig. 5B). Thus, the presence of *fyuA* in AIEC significantly altered host transcriptional responses in the inflamed colon prior to and throughout the development of fibrosis.

**Figure 5.**
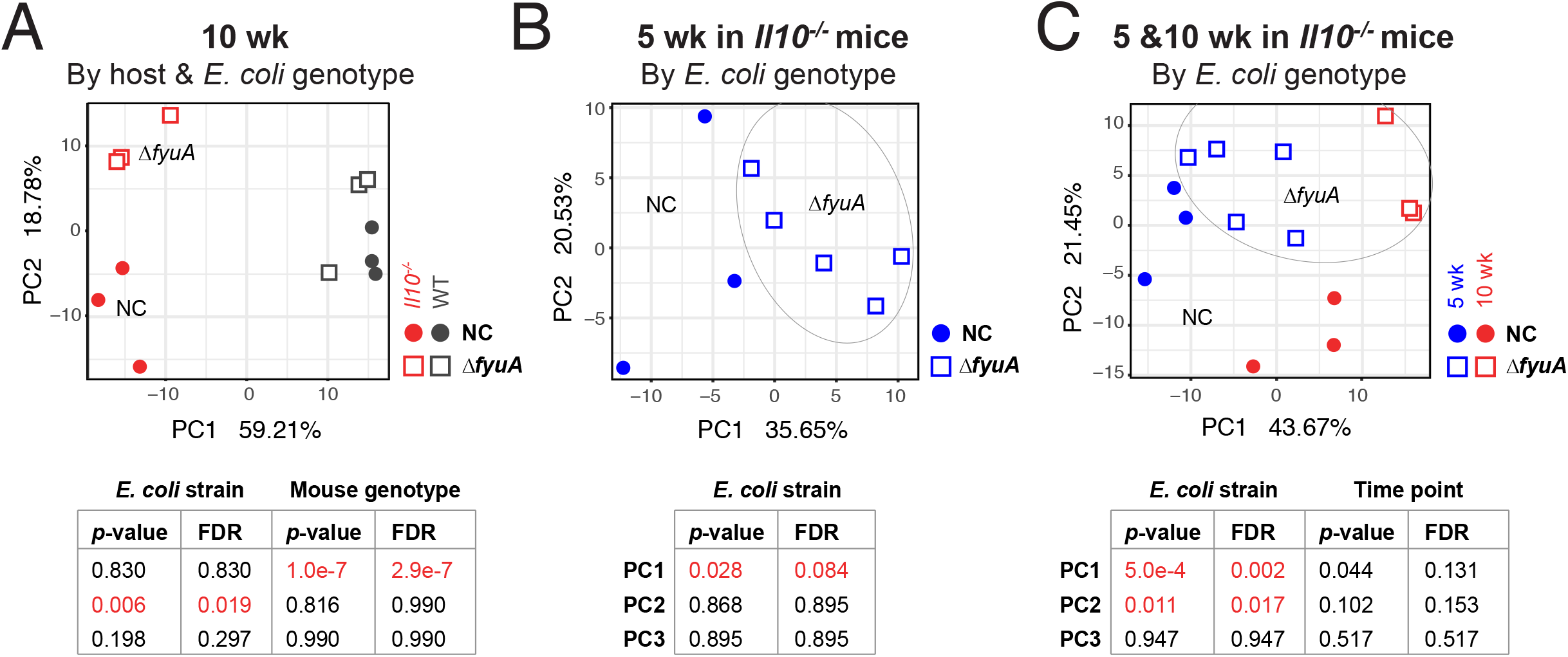
Deletion of *fyuA* in AIEC promotes transcriptome-wide changes in the colons of *Il10^−/−^* mice. Principal component analysis of transcriptome-wide changes in the colons of A) NC-vs Δ*fyuA*-colonized WT or *Il10^−/−^* mice after 10 weeks, B) NC-vs Δ*fyuA*-colonized *Il10^−/−^* mice after 5 weeks or C) NC-vs Δ*fyuA*-colonized *Il10^−/−^* mice after 5 or 10 weeks.

Transcriptomic analysis of a prospectively followed inception cohort of pediatric CD patients revealed high expression of profibrogentic genes and pathways prior to the development of stricturing fibrotic disease (30). This included ECM structural constituents and collagen binding pathways (30). In agreement with these results, the ECM-receptor interaction KEGG pathway is significantly upregulated in Δ*fyuA*-colonized mice during (10 weeks) and prior to (5 weeks) histological evidence of fibrosis (Table S2, S4). We generated a heat map to visualize expression of individual genes in this KEGG pathway between individual NC101-versus Δ*fyuA-* colonized *Il10^−/−^* mice (Fig. 6A). Phylogenetic clustering of the 5-week samples demonstrated that three of the Δ*fyuA*-colonized mice clustered together and exhibited increased expression of numerous ECM genes, including type I, IV and VI collagens and fibronectin (arrowheads, Fig. 6A). Careful histological observation by a pathologist blinded to the treatment groups revealed early evidence of fibrosis in these three Δ*fyuA-* colonized mice, but not in the remaining Δ*fyuA*-colonized mice that clustered with the NC101-colonized mice and exhibited lower expression of ECM genes. These unbiased molecular findings are consistent with our observation that a subset, and not 100%, of Δ*fyuA*-colonized mice develop fibrosis. These findings were confirmed by targeted quantitative PCR analysis, where transcript levels of *col1a2* (type 1 collagen) and *fn1* (fibronectin) were significantly increased in *fyuA-* vs NC101-colonized *Il10^−/−^* mice (Fig. 6B-C). This corresponded with increased positivity of α-SMA (smooth muscle actin), a common feature of fibrosis, in Δ*fyuA*-colonized *Il10^−/−^* mice. Taken together, these findings demonstrate that ECM components are upregulated in pre-fibrotic *Il10^−/−^* mice colonized with Δ*fyuA* prior to the development of fibrotic disease.

**Figure 6.**
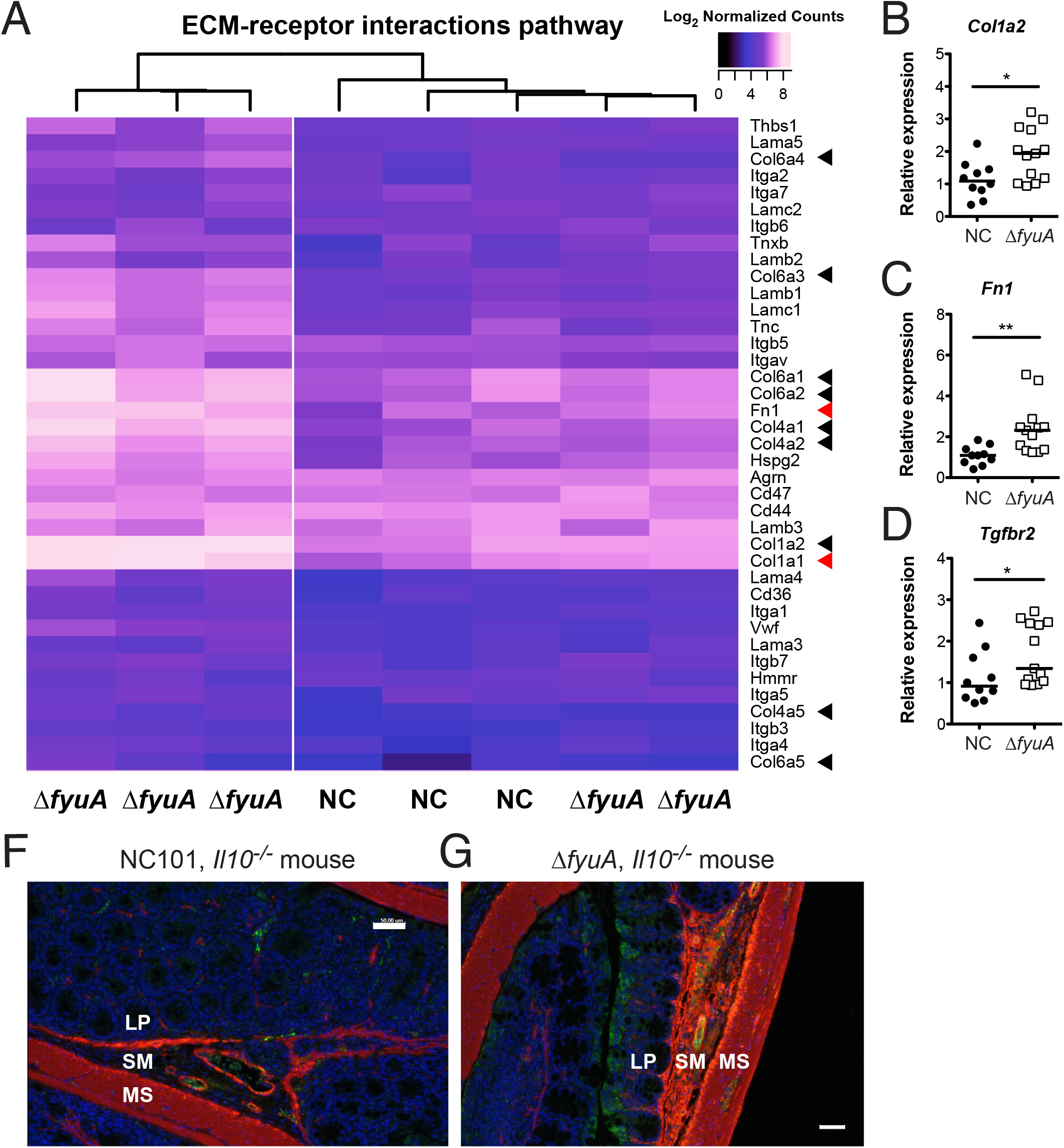
Deletion of *fyuA* in AIEC promotes pro-fibrotic host responses preceding fibrosis development. A) Heat map of log2 normalized counts of genes in the ECM-receptor interaction KEGG pathway in NC- or Δ*fyuA*-colonized *Il10^−/−^* mice at 5 weeks. B-D) Relative colonic transcript levels of B) *Col1a1*, C) *Fn1* and D) *Tgfbr2* in *Il10^−/−^* mice mono-associated with NC or Δ*fyuA* for 5 weeks. Each symbol represents an individual mouse (n = 10-13). Lines are at the median. *P*-values were determined by Mann-Whitney. * *p* < 0.05, ** *p* < 0.01. F-G) Proximal colons from *Il10^−/−^* mice colonized with F) NC or G) Δ*fyuA* for 10 weeks were stained with α-SMA (red), CD31 (green), or DAPI. LP, lamina propria. SM, submucosa. MS, muscularis serosa. Scale bar, 50 μm.

TGF-β signaling represents the canonical pro-fibrotic activation pathway. Therefore, to further confirm the presence of a pro-fibrotic gene signature in fibrotic mice, we evaluated the expression of genes within the TGF-β pathway. RNAseq analysis detected colonic expression of the three TGF-β and TGF-β receptor isoforms. Pre-fibrotic (5 weeks) Δ*fyuA-* versus NC101-colonized *Il10^−/−^* mice trended towards elevated expression of TGF-β1 and TGF-β3 and TGF-β receptor isoforms 1 and 2 (Fig. S10A). Increased expression of the TGF-β2 receptor was confirmed by quantitative PCR (Fig. 6D). Similarly, a significant increase in TGF-β 1-3 and TGF-β receptor isoforms 2 and 3 expression was observed at 10 weeks in fibrotic Δ*fyuA*-colonized *Il10^−/−^* mice (Fig. S10b). Together, these data further support our hypothesis that deletion of *fyuA* in AIEC promotes a profibrogenic environment in inflammation-susceptible hosts, which occurs at an early phase of the inflammatory response prior to onset of fibrosis.

### Ybt-dependent fibrosis is not associated with altered host systemic iron homeostasis

Membrane permeable siderophores like Ybt disrupt host iron homeostasis and modulate iron sensitive host responses, which includes the induction of *Ndrg1* (32) (33). Because deletion of *fyuA* does not alter Ybt secretion, colonization with Δ*fyuA* may instead increase Ybt internalization by host cells in the absence of bacterial import and alter host iron homeostasis to promote fibrosis. To address this possibility, we determined whether colonization with Δ*fyuA* versus NC101 or Δ*irp1* alters systemic iron homeostasis in *Il10^−/−^* mice. At 2 weeks (prior to histological inflammation or fibrosis) and at 10 weeks (when colitis and fibrosis are evident in affected animals), plasma hemoglobin levels did not differ (Fig S11A-B). Similarly, Prussian blue staining did not reveal differences in splenic iron stores at 10 weeks (Fig S11C). To determine whether local iron homeostasis was altered in the colon, we utilized our RNAseq data to assess whether established host iron-responsive genes were differentially expressed in mice colonized with NC101 or Δ*fyuA* (Table S5) (34) (35) (36) (37). Of the 15 canonical iron-responsive genes investigated, three were differentially regulated in *Il10^−/−^* mice including *Ndrg1* and *Tfrc* (transferrin receptor) and two were differentially regulated in WT mice including *Tfrc* at 10 weeks. At 5 weeks, *Epas1* was the only iron-responsive gene that was altered between NC101-versus Δ*fyuA*-colonized *Il10^−/−^* mice, a change not observed at 10 weeks. Together, these findings suggest that Δ*fyuA* does not profoundly alter systemic or colonic iron homeostasis in the host and may not be a driving factor for fibrosis induction.

### Yersiniabactin biosynthesis is required for AIEC-mediated fibrosis induction

Abrogation of Ybt transport in AIEC had opposing effects on fibrosis induction in *Il10^−/−^* mice compared to the inactivation of Ybt biosynthesis (Fig. 2). Because fibrosis development was minimal in *Il10^−/−^* mice colonized with the Δ*irp1* mutant, we next determined whether Ybt biosynthesis is required for the fibrosis-inducing potential of Δ*fyuA*. Genetic inactivation of Ybt biosynthesis in Δ*fyuA* (Δ*fyuAirp1*) significantly reduced fibrosis incidence in *Il10^−/−^* mice (Fig. 7A-C). Moreover, when comparing fibrosis incidence in mice colonized with Ybt-positive versus Ybt-negative AIEC, 22 out of 51 mice colonized with Ybt-positive AIEC developed fibrotic disease, whereas 3 out of 26 mice colonized with Ybt-deficient AIEC exhibited histological evidence of fibrosis (Fig. 7B). Inactivation of Ybt production in Δ*fyuA* also reduced its subepithelial invasiveness, resulting in a similar pattern of tissue localization compared to NC101 (Fig. 7D). This further reinforces the link between increased mucosal invasiveness and the enhanced profibrogenic potential of Δ*fyuA*. Importantly, colitis severity at 10 weeks was comparable between Δ*fyuA* and Δ*fyuAirp1* (Fig. S12), suggesting that differences in inflammation were not driving fibrosis severity.

**Figure 7.**
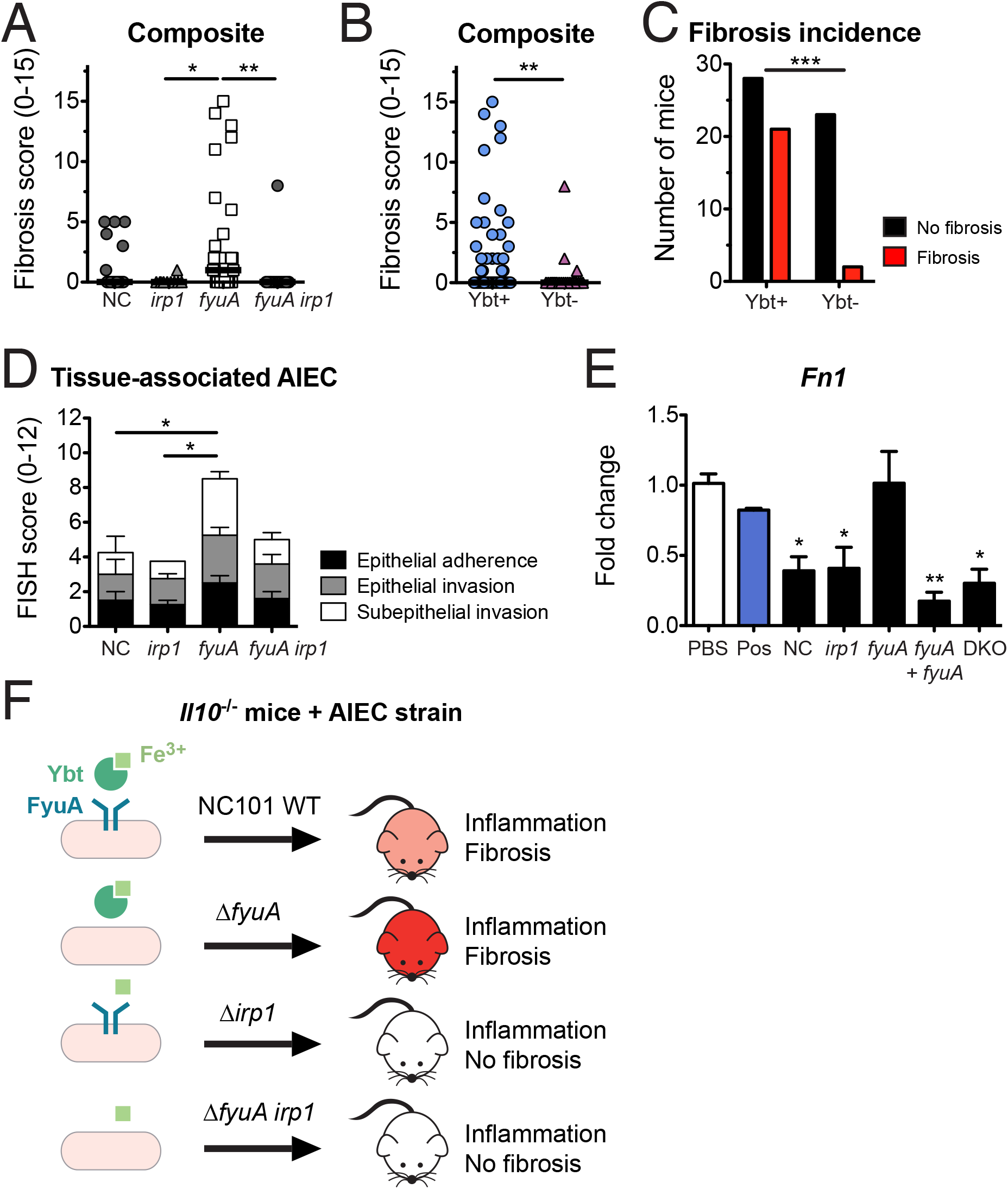
Yersiniabactin biosynthesis promotes fibrosis in AIEC-driven colitis. Germ free *Il10*^−/−^ mice were mono-associated with the AIEC strain NC, Δ*fyuA, Δirp1* or Δ*fyuA irp1* for 10 weeks. A) Composite fibrosis histology scores. Each symbol represents an individual mouse (n = 10-29). Lines are at the median. *P*-values were determined by Kruskal-Wallis. B) Composite histopathology colitis scores of *Il10^−/−^* mice colonized with ybt-positive or ybt-deficient AIEC. Lines are at the median. *P*-values were determined by Mann-Whitney. C) Fibrosis incidence rates of *Il10^−/−^* mice colonized with Ybt-positive or Ybt-deficient AIEC as assessed by H&E histology. *P*-values were determined by Fisher’s exact test. D) FISH analysis of proximal colons (*n* = 4-8). *P*-values were determined by one-way ANOVA. E) Swiss 3T3 fibroblasts were co-cultured with NC, Δ*irp1, ΔfyuA, ΔfyuA+fyuA* or Δ*fyuA irp1*. Fibroblasts stimulated with TGF-β served as a positive control (pos). Data are represented as the mean ± SEM. *P*-values were determined by Kruskal-Wallis. F) Working model. Data for NC-, Δ*fyuA*-, and Δ*irp1-* colonized mice are also presented in Figures 2 and 4. * *p* < 0.05, ** *p* < 0.01, *** *p* < 0.001.

To further demonstrate the profibrogenic potential of Δ*fyuA*, we next determined whether inactivation of Ybt transport in AIEC enhances the activation of cultured fibroblasts *in vitro*. Fibroblasts that were cultured with Δ*fyuA* expressed significantly higher levels of the fibroblast activation marker *Fn1* in comparison to the parental and Ybt-deficient strains (Fig. 7E). This corresponded with our *in vivo* observations, where *Fn1* transcripts were elevated in Δ*fyuA*-colonized *Il10^−/−^* mice (Fig. 6C). Together, these results demonstrate that inactivation of the Ybt siderophore system in AIEC in two distinct manners (i.e. Ybt transport versus Ybt biosynthesis) does not have similar effects on colitis induction and fibrosis development in genetically susceptible hosts. More broadly, in addition to its role in bacterial iron acquisition, our findings collectively introduce a novel, non-canonical role of Ybt in establishing a profibrogenic microenvironment in inflammation-susceptible hosts.

## Discussion

Siderophore biosynthetic gene clusters are abundant in the gut microbiota, with 232 putative clusters identified from metagenomes in the Human Microbiome Project study (38). Given that IBD-associated AIEC strains also harbor many of these siderophore systems (21), it is conceivable their siderophores may contribute to AIEC-associated intestinal disease. Indeed, here we introduce the siderophore Ybt as a novel bacterial factor that promotes profibrogenic host responses in the inflamed intestinal environment. Our findings demonstrate that AIEC are pro-fibrogenic, and inactivation of Ybt transport in a colitogenic AIEC strain enhances fibrosis development in inflammation susceptible mice. Inactivation of Ybt biosynthesis in both the Ybt transport mutant and the parental strain abrogates their fibrosis-inducing potential, suggesting that Ybt promotes fibrosis development even in the absence of uptake through its canonical receptor. Profibrogenic transcriptional signatures are evident in the colon prior to histological presentation of disease, suggesting a causative role for Ybt-mediated induction of fibrosis. Together, our findings introduce a specific microbiota-derived factor that promotes the development of inflammation-associated fibrosis.

The canonical function of the Ybt siderophore system is to import extracellular iron sequestered by Ybt through FyuA for bacterial use. Thus, inactivation of FyuA may enhance the profibrogenic potential of AIEC by perturbing bacterial iron homeostasis and subsequently modulating bacterial function. However, luminal expression of highly-sensitive, iron-responsive genes (39) (31) were comparable between the NC101 parental strain, the transport mutant Δ*fyuA* and the Ybt-deficient mutant Δ*irp1*. This indicates a lack of strain-specific differences in iron sensing. Similarly, while the functional outcome of iron starvation in bacteria is a fitness disadvantage (40) (31), we observed no prolonged differences in luminal colonization between NC101, Δ*fyuA* or Δ*irp1* in the non-inflamed intestines or during the course of inflammation and fibrosis development. This is likely the result of additional iron scavenging systems in NC101 that serve compensatory roles in the Ybt-deficient and transport mutants (41) (21).

Indeed, in other Enterobacteriaceae strains, inactivation of multiple iron acquisition systems is required to attenuate *in vivo* fitness (25) (31). Most importantly, if the enhanced profibrogenic potential of Δ*fyuA* was the result of dysregulated bacterial iron homeostasis and consequent effects on Ybt-independent functions, we would expect fibrosis induction mediated by Δ*irp1* and Δ*fyuA* to be comparable, as both mutants cannot scavenge iron through Ybt (42). Instead, fibrosis induction was further attenuated in mice colonized with Ybt-deficient AIEC strains. Collectively, our findings support a model where Ybt stimulates host profibrogenic responses through a mechanism independent of its role in importing iron through FyuA.

While the Ybt system did not impact overall AIEC intestinal fitness, inactivation of Ybt transport altered the distribution of AIEC colonization within colonic tissues. This may contribute to AIEC-driven fibrosis by activating myofibroblasts and mesenchymal cells either directly via bacterial recognition receptors (i.e. TLRs) or indirectly by activating intestinal immune cells that modulate profibrogenic cellular responses (5) (6). In comparison to the parental strain, Δ*fyuA* was more abundant within the colonic subepithelium and co-localized with fibrotic lesions in *Il10^−/−^* mice at 10 weeks. In contrast, inactivation of Ybt biosynthesis did not alter tissue localization of AIEC, further uncoupling the effects of Ybt biosynthesis and Ybt transport on bacterial function. Instead, inactivation of Ybt biosynthesis in Δ*fyuA* restored tissue colonization patterns exhibited by the parental strain, suggesting that Ybt mediates the mislocalization of Δ*fyuA* to the subepithelium independent of its role in importing iron through FyuA. Consistent with our findings, several studies have also reported altered tissue localization of Enterobacteriaceae pathogens with inactivation of the Ybt system in extraintestinal mucosal environments (43) (26) (44). Finally, it should be noted that while we observed a statistically significant decrease in fecal colonization of the Δ*fyuA* mutant at 5 weeks, it remains unclear whether a <0.5 log difference in bacterial burdens in a mono-colonized mouse can impart any meaningful effects on the host – especially as this decrease was also not observed at 1 or 10 weeks post colonization. Collectively, these findings highlight one putative non-canonical function of Ybt that may enhance the profibrogenic potential of AIEC. The precise mechanisms by which inactivation of FyuA enhances fibrosis development in susceptible hosts will be the subject of future studies.

Because Ybt is a secreted bacterial product that permeates mammalian membranes, Ybt may also promote profibrogenic host responses by perturbing cellular iron homeostasis in the host. Indeed, the membrane permeable siderophores enterobactin and yersiniabactin stimulate epithelial proinflammatory responses by decreasing intracellular iron pools, an effect that is reversed with the addition of iron (32) (33). Ybt disruption of local iron homeostasis may similarly drive fibrosis development by stimulating profibrogenic responses in epithelial, mesenchymal and immune cells. While our host transcriptomics analyses demonstrated similar colonic expression profiles of numerous iron responsive host genes in NC101-versus Δ*fyuA*-colonized mice (Table S5), differences in the canonical iron-response genes *Ndrg1* and *Tfrc* were uniquely observed in *Il10^−/−^* mice. This suggests that local and/or cell-specific alterations in host iron homeostasis may contribute to the progression of fibrosis. However, as these changes were observed at 10 weeks but not 5 weeks post-colonization, the initiation of fibrosis and early pro-fibrotic gene signatures cannot be attributed to major alterations in host iron homeostasis.

In addition to iron, Ybt is capable of binding other metals including nickel, cobalt, chromium, gallium, and copper (45). This raises the possibility that its profibrogenic potential is the result of interactions with other metals present in the colonic environment. For example, when complexed with copper, Ybt acts to limit the lethal effects of macrophage reactive oxygen species (46). Ybt-copper chelates have been detected in urine samples from patients infected with uropathogenic *E. coli* (UPEC), demonstrating that Ybt binds copper *in vivo* (23). Bacterial cells can also import Ybt-copper chelates through FyuA (47) (24). Together, these findings introduce two putative mechanisms by which Ybt promotes fibrosis development: 1) through chelation of host sources of metals other than iron and/or 2) by modulating the transcriptome and metabolome of AIEC by altering the flux of micronutrients into the bacterial cell. Finally, because the Ybt enzymatic machinery produces additional secreted metabolites that remain uncharacterized (48), it is intriguing to speculate that these Ybt precursors and Ybt-like molecules may also play a role in inflammation-associated fibrosis.

Fibrosis complicates many inflammatory intestinal disorders associated with microbial dysbiosis, however, pro-fibrotic mechanisms remain incompletely understood and limit therapeutic strategies. This has been hampered by the lack of rodent models that recapitulate the complex interactions between host genetics and microbial factors important for inflammation and fibrosis development. Here, we introduce a new model for inflammation-associated fibrosis driven by a pathobiont-derived small molecule produced from the Ybt pathogenicity island. Consistent with human CD (30), profibrogenic pathways are upregulated prior to histological presentation of fibrosis and mirror the incidence rate of fibrotic disease in our model. Moreover, our model recapitulates key histological and transcriptomic aspects of fibrotic disease in human CD. More broadly, our findings demonstrate that manipulating the same pathogenicity island in different ways can result in distinct consequences for disease development. This highlights an important difference in targeting siderophore biosynthesis versus the cognate receptors as putative bacterial targets in microbial driven diseases such as CD. Furthermore, other siderophore and metallophore systems of the gut microbiota may induce similar responses and contribute to fibrosis. Given the prevalence of AIEC among the CD population, the presence of the Ybt siderophore system could serve as a useful prognostic tool in identifying patient subsets susceptible to fibrotic disease.

## Materials and methods

### Bacterial strains

The fecal isolate *E. coli* NC101 was isolated from WT mice (13). The *λ-red* recombinase system was utilized to generate mutants (49) (Table S7). Bacterial strains and plasmids are listed in Table S6.

### Mice

Germ free *Il10^−/−^* and WT 129S6/SvEV mice were maintained at the National Gnotobiotic Rodent Resource Center at UNC-CH. Absence of isolator contamination was confirmed by Gram stain and fecal culture. Eight-12-week old mice were inoculated via oral and rectal swab with *E. coli* following overnight growth in LB broth (13). Colonization was confirmed by fecal plating. Five cohorts of *Il10^−/−^* mice were colonized with NC101 WT or Δ*fyuA* and two cohorts of *Il10^−/−^* mice were colonized with Δ*irp1* or Δ*fyuAirp1*. Animal protocols were approved by the UNC-CH Institutional Animal Care and Use Committee.

### Quantification of bacteria

*E. coli* CFUs in feces were quantified by serial dilutions and plating on LB plates. Mucus- and tissue-associated bacteria were enumerated as described (10).

### Colitis histopathology

At necropsy, tissues were fixed in 10% neutral buffered formalin. Colon sections were stained with H&E, Masson’s trichrome or Sirius red; spleens with Prussian blue. Colitis scores (0-12) of Swiss-rolled colons were blindly assessed as described (13) (20). Composite scores (0-36) are the sum of proximal, middle and distal colon scores.

### Fibrosis histopathology

Fibrosis was blindly assessed on colonic H&E sections and validated by Sirius red. Severity of fibrosis (0-5) was evaluated using a validated scoring system (28) (29) evaluating the extent of submucosal involvement: 0 – no fibrosis; fibrosis in: 1 - <25%, 2 - 26-50%, 3 - 51-75%, or 4 - 76-100% of colon section. One point was added for lamina propria involvement. Composite scores (0-15) are the sum of the proximal, middle and distal colon scores. 0 was considered non-fibrotic, 1-3 represented mild fibrosis, and 4+ represented moderate/severe fibrosis. Histopathology was blindly confirmed by a small animal veterinarian specializing in gastrointestinal histopathology.

### Human intestinal samples

Formalin-fixed, paraffin embedded tissue blocks from routine diagnostic surgical resections were transferred to UNC-CH under approved Institutional Review Board protocol of the Cleveland Clinic. Sections were H&E and Sirius red stained from three individuals per disease category: CD, UC, diverticulitis, and non-inflamed controls (healthy margins of colorectal cancer patients).

### Fluorescent *in situ* hybridization (FISH)

Colons were washed in PBS to remove contents and loosely adhered bacteria. Formalin-fixed, paraffin-embedded sections were mounted on charged glass slides and incubated with an oligonucleotide probe directed against *E. coli* (Cy3-E. *coli/Shigella* probe) and an antisense probe (6-FAM-non-EUB338) (50). Hybridized samples were washed in PBS, air dried, and mounted with ProLong antifade (Molecular Probes Inc.). Sections were examined on a BX51 epifluorescence microscope with Olympus DP-7 camera. The FISH analysis was performed in a blind fashion by two independent investigators as follows: to assess bacterial colonization, we enumerated individual bacterial cells adhered to epithelial cells (epithelial attachment), localized within epithelial cells (epithelial invasion), and translocated across the epithelium (subepithelial invasion). The quantity of bacteria per colon swiss-roll was converted to a FISH score of 0-4 (Table 1).

**Table 1:** FISH Scoring

| FISH SCORE | Epithelial attachment | Epithelial invasion | Subepithelial invasion |
| --- | --- | --- | --- |
| 0 | None | None | None |
| 1 | 1-50 | 1-10 | 1-5 |
| 2 | 51-150 | 11-20 | 6-10 |
| 3 | 151-250 | 21-30 | 11-15 |
| 4 | 251+ | 31+ | 16+ |

### RNA-seq analysis

RNA-seq reads were quality filtered at Q20 and trimmed to remove remaining adaptors using Trimmomatic (51) version 0.35. Resulting reads were aligned to Illumina iGenome *Mus musculus* GRCm38 reference genome using Tophat (52) version 2.1.0 utilizing Bowtie2 (53) version 2.2.5. Resulting alignments were processed using Cufflinks (54) version 2.2.1 along with Illumina iGenome *Mus musculus* GRCm38 Gene transfer format file, after masking rRNA features as described (55). Transcripts were quantified using cuffquant and gene counts were exported to text files and then imported to edger (56) version 3.12.1 (running inside R version 3.2.3) for detecting differentially expressed genes. A gene was considered for differential expression test if present in at least three samples. We considered a gene differentially expressed if its edgeR FDR adjusted p-value < 0.05. Parallel analysis using featureCounts (57) from the subread package version 1.4.6 for transcript quantification showed similar results.

Pathway analysis was conducted using GAGE (58) version 2.20.1 using *Mus musculus* (mmu) Kyoto Encyclopedia of Genes and Genomes (KEGG) (59) pathways and genes were mapped to KEGG pathways using Pathview version 1.10.1 (60). Pathways were considered significant if its GAGE q-value was <0.05. ECM-receptor interactions pathway genes (Figure 6) are based on KEGG pathway mmu04512. We tested the effect of sequencing run and lane on the clustering of the samples and found both to be insignificant (P-value > 0.05) for PC1 and PC2 in Figure 5.

### Statistical analysis

*P*-values were calculated using non-parametric Mann-Whitney when 2 experimental groups were compared or Kruskal-Wallis with Dunn’s post-test when ≥3 experimental groups were compared. Data from quantitative bacterial culture were log transformed for normalization. *P*-values <0.05 were considered significant.

Additional methods are described in the Supplemental Materials and Methods.

## Acknowledgements

We acknowledge the Histology and Gnotobiotic Cores at the UNC Center for Gastrointestinal Biology and Disease (CGIBD: supported by NIH P30DK34987), the Translational Pathology Laboratory at the UNC Lineberger Comprehensive Cancer Center (supported by NCI (P30-CA016086-40), NIEHS (P30ES010126-15A1), UCRF, and NCBT (2015-IDG-1007)) and the National Gnotobiotic Rodent Resource Center (supported by NIH P40 OD01995). We thank the University of North Carolina’s Department of Chemistry Mass Spectrometry Core Laboratory, especially Brandie Ehrmann, for their assistance with mass spectrometry analysis, supported by NSF (CHE-1726291) and the UNC-CH School of Medicine Office of Research. We also acknowledge Taylor Tibbs and Lacey Lopez for assistance with histological stains, and Elise Sloey for assistance with GFP reporter assays. This work was supported by the following grants: NIH/NIDDK K01 DK103952 (JCA), Kenneth Rainin Foundation Innovator Award (JCA), Pilot feasibility grant from the CGIBD to JCA (NIH P30DK34987), American Gastroenterological Association Augustyn Award in Digestive Cancer (JCA), Lineberger Comprehensive Cancer Center Pilot Grant and UCRF (JCA), North Carolina Translational and Clinical Sciences Institute (NC TraCS) pilot grant (JCA), American Cancer Society postdoctoral fellowship (JCA), UNC dissertation completion fellowship (ME), Microbiome Initiative (RBS and KWS) and Gnotobiotic Facility (RBS) grants from the Crohn’s and Colitis Foundation of America (CCFA), a Student Research Fellowship Award from the CCFA (LNM), and pilot grant P30DK097948 and K08DK110415 (FR).

## Author contributions

Conceptualization, M.E., K.S., R.B.S. and J.C.A.; Methodology, M.E. and J.C.A.; Software, R.Z.G.; Validation, A.R.; Formal Analysis, M.E., R.Z.G., B.D. and J.C.A.; Investigation, M.E., L.F., B.D., L.N.M, C. A. B, L.R.L., A.M.R., C.B., J.W.H, I.O.G., and J.C.A.; Resources, C.B., K.S., C.J., I.O.G., F.R., R.B.S. and J.C.A.; Data Curation, R.Z.G.; Writing – Original Draft, M.E. and J.C.A.; Writing – Review and Editing, M.E., J.C.A and R.B.S.; Visualization, M.E.; Supervision, J.C.A.; Funding Acquisition, J.C.A. and R.B.S.

